# Prenatal circadian rhythm disruption induces sex-specific substance use and mood-related phenotypes in mice

**DOI:** 10.64898/2026.06.22.733807

**Authors:** Nilanjana Saferin, Taylor Stowe, Chelsea A. Vadnie, Kaitlyn A. Petersen, Madeline R. Scott, Emily Chen, Lucía Bustos-Robles, Riley Griffin, Colleen A. McClung, Lauren M. DePoy

## Abstract

20% of Americans are at risk for environmental circadian rhythm disruptions (CRD) due to shift work, leading to substantial negative health outcomes. However, females are especially affected with greater vulnerability for substance use (SU) and adverse outcomes associated with pregnancy, including for offspring at birth and later in life. In mice, prenatal CRD (pCRD) recapitulates these risks, but it is unknown whether pCRD affects SU in mature offspring. To investigate this, C57BL/6J dams were disrupted by reversing the light/dark cycle during gestation. Following pCRD, reward- (cocaine conditioned place preference, intravenous self-administration) and mood-related behaviors (open field, elevated plus maze, light/dark box, forced swim) were measured in adult offspring. Adult female offspring of dams exposed to CRD developed an anhedonic-like phenotype with decreased food self-administration, cocaine intake and reinforcing properties of cocaine. Opposingly, pCRD male offspring showed a SU-like phenotype with increased cocaine preference, higher order food self-administration and cocaine reinforcement. Interestingly, these divergent behavioral outcomes were not specific to reward. While female pCRD mice showed increased anxiety-like behavior, pCRD males showed decreased anxiety/increased risk-taking behavior, as well as decreased immobility in the forced swim test. Rhythms in corticosterone were also sex-specifically affected by pCRD. These results suggest that pCRD may predispose individuals to distinct psychiatric disorders based on sex with mood disorders developing in females and SU disorders developing in males. By better understanding how disrupted rhythms during pregnancy affect behavior in adulthood, we can develop novel therapeutic approaches for SU and mood disorders in adults.

## Introduction

Psychiatric disorders and circadian rhythm disruptions (CRD) often co-occur, with circadian misalignment, delayed rhythms, sleep disruptions and social jet lag frequently seen in conditions including depression^1–3^ and substance use disorders (SUD)^4–7^. SUDs, in particular, represent a major public health concern, affecting 1.5 million Americans, with psychostimulant overdoses doubling in recent years and no current FDA-approved treatments available^8^. A bidirectional relationship exists, where drug use disrupts circadian rhythms^9–12^, and individuals with disrupted sleep or circadian rhythms are more susceptible to drug use^13–16^. Environmental and genetic disruptions can perturb circadian rhythms by impacting the molecular clock, a series of interlocking transcriptional-translational feedback loops found in almost every cell in the body^17–19^, which controls physiology and behavior across the light/dark (LD) cycle. The molecular clock is largely regulated by a dimer of CLOCK (circadian locomotor output cycles kaput) and BMAL1 (brain and muscle ARNT-like protein 1), which control transcription of many genes including Period (*Per*) and Cryptochrome (*Cry*)^18^. After translation, the PER and CRY proteins enter the nucleus and inhibit their own transcription, closing the negative feedback loop after around 24□hours^17^. Both environmental and genetic disruptions to circadian rhythms are linked to altered mood^20–23^ and substance use (SU)^24–30^, making CRD a key vulnerability factor in identifying at-risk populations and developing therapeutic strategies to treat SUD.

Shift workers, currently 20% of the U.S. workforce, are the largest at-risk group for severe environmental CRD^31^, which increases their susceptibility to metabolic, cardiovascular, gastrointestinal^32^, mood disorders^33^, and SU^34^. Female shift workers are particularly vulnerable, experiencing higher rates of SU and depression^33,37^, as well as pregnancy complications such as preterm birth and low infant weight, with risks intensifying as CRD severity increases^38^. Rodent models of CRD exhibit comparable negative health effects, including metabolic and cardiac dysfunction^35,36^. Although research on the effect of CRD on SU in mice is limited, studies have reported increased alcohol consumption and altered drinking patterns in adult animals^39,40^. Furthermore, in rodents, prenatal CRD (pCRD) reduces full-term pregnancy rates^41^, supporting light/dark cycle shifting in mice as a relevant model for investigating pCRD in humans.

The long-term effects of pCRD on offspring remain unclear, but studies link maternal CRD severity to increased depression risk in children^42^. Notably, children of morning chronotype (i.e. people with a preference for morning activity) shift workers, who experience greater circadian misalignment, show higher depression rates than those of evening chronotypes, suggesting the severity of circadian disruption, not shift work itself, drives these outcomes^42^. Adolescents with shift-working parents also exhibit more risk-taking^43^ and SU^44^, but separating prenatal from postnatal influences in human studies is challenging. In mice, pCRD increases depression- and anxiety-like behaviors^45–47^, though its effect on reward-related behaviors is unknown. Understanding the link between shift work and SU in parents and offspring, as well as the long-term risks of pCRD is essential. Here, we sought to investigate the long-term effects of pCRD by exposing mice to 12-h shifts in the LD cycle during gestation. This type of LD cycle shifting is known to reduce novelty seeking in adult mice^48^, as well as shift the phase and dampen the amplitude of activity rhythms^36^, disruptions that are also implicated in psychiatric disorders in humans^1,49,50^. In this study, we hypothesized that pCRD would have persistent effects on reward- and mood-related behavior in adult offspring. By understanding how disrupted rhythms during pregnancy affect behavior in adulthood, we can develop novel therapeutic approaches for SU and mood disorders in adults.

## Materials and Methods

### Animals and housing conditions

Male and female C57BL/6J mice were housed under a 12-hour (h) light/dark (LD) cycle, with lights turning on at 0700 (Zeitgeber Time, ZT0), except during circadian disruption. Behavioral assessments were conducted during the light phase (ZT2-7), unless otherwise noted. Food and water were provided ad libitum unless otherwise indicated. All procedures were approved by the Institutional Animal Care and Use Committee.

### Circadian rhythm disruption

CRD was initiated in 8–12-week-old pregnant dams. Three days after harem breeding pairs were established, dams were separated and exposed to 12-h shifts every 5 days during gestation, resulting in 4 complete reversals of the LD cycle before birth. Control animals were sham handled and kept on a normal schedule. Offspring were born under a normal LD cycle (ZT0 = 0700). Behavioral tests were performed in adult offspring (8-20 weeks of age). Pup mortality and body weight, litter size and pup retrieval (∼P7) were measured. Homecage activity and sleep/wake measurements were also recorded for a subset of dams during gestation (PiezoSleep, Signal Solutions, LLC, Lexington, KY, United States).

### Drugs

Cocaine hydrochloride was obtained from the National Institute on Drug Abuse. For conditioned place preference testing, animals received intraperitoneal (i.p.) injections of 5 mg/kg (volume: 10 mL/kg). For cocaine self-administration, doses ranged from 0 to 1 mg/kg per infusion.

### Behavioral testing

Mice used for behavioral testing were assigned to only one of five different cohorts, which are described briefly. Further details for each behavioral test can be found in the supplement. All behavior was conducted during the light phase at ∼ZT2, except for cohort 3, which were trained during both the light phase (∼ZT2) and dark phase (∼ZT14).

#### Cohort 1: Exploratory drive

Mice underwent a battery of tests in the following order to evaluate locomotor activity and exploratory behavior using open field, elevated plus maze (EPM), and light/dark (LD) box as well as depression-like behavior (forced swim test, FST). Testing was conducted every other day. Mice received the tests in this order, but some cohorts received a subset of tests (*e.g.* not all mice were tested in the FST).

#### Cohort 2: Reward-related behaviors

Mice were assessed for reward-related behavior through sucrose preference and cocaine-conditioned place preference (CPP)^51^ tests.

#### Cohort 3: Cocaine self-administration

Mice were trained on various schedules of reinforcement for food self-administration and then intravenous cocaine self-administration. This included a fixed ratio (FR1) schedule for food and drug self-administration, followed by dose response curve, progressive ratio, extinction, and cue-induced reinstatement for drug self-administration.

#### Cohort 4: Food self-administration

Mice were trained to lever press for food reinforcers at various schedules of reinforcement.

#### Cohort 5: Action-outcome contingency degradation

Mice were trained to lever press two levers for food reinforcers at various schedules of reinforcement. Next, action outcome contingency degradation was assessed.

#### Cohort 6: Cross fostering

Pups from a control dam were fostered by a pCRD dam and vice versa. Mice in this cohort were evaluated for a subset of exploratory behavior such as open field and elevated plus maze, and reward-related behaviors such as conditioned place preference.

### Blood collection

We measured rhythms in corticosterone (CORT), as well as the glucocorticoid response to acute restraint stress^52^. For CORT rhythms, adult mice (≥12 weeks old) exposed to pCRD and controls were euthanized across time of day (TOD)(ZT2/7/10/14/19/22), and trunk blood was collected for plasma CORT measurements. For acute restraint stress, adult pCRD and control mice were restrained in 50-mL conical tubes between ZT2 and ZT7. Tail blood samples were collected at baseline, after 15 min restraint, as well as at 30-and 120-minutes. Plasma was isolated and stored at −80 °C until analysis.

### Corticosterone ELISA

Plasma corticosterone (CORT) samples were diluted 1:100 and CORT levels were quantified as instructed in a commercially available ELISA kit (ab108821, Abcam). Absorbance was determined at a wavelength of 450 nm and CORT concentration was determined based on the standard curve. All samples were run in triplicate and mean values were used for data analysis.

### Statistical Analyses

GraphPad Prism 10 and SPPS were used. Data are expressed as mean ± SEM with *p* ≤ 0.05 considered significant and 0.05 < *p* ≤ 0.1 considered trending. Measures collected in CRD dams and litters (e.g. sleep, pup mortality) were analyzed by two-tailed t-tests. Homecage activity and sleep were analyzed in dams using WakeActive/ActivityStatistics and SleepStats2p10 software respectively (Signal Solutions, LLC, Lexington, KY, United States).

For measures collected in adult offspring, ANOVAs were performed with repeated measures (RM) as needed. Trending and significant interactions were followed by Bonferroni post-hoc tests corrected for multiple comparisons. For sucrose preference, CPP and action outcome contingency degradation, one sample t-tests were used to determine whether significant preferences exist (compared to 50%, 0 and 1 respectively).

Additional details are found in the supplemental material.

## Results

Pregnant mice were exposed to CRD, consisting of four 12-h shifts in the LD cycle occurring every 5 days during gestation (Fig.1A) and behavior was tested in adult offspring (>8 weeks old). To ensure our model of CRD disrupted rhythms during pregnancy, we assessed locomotor activity and sleep in pregnant dams. Actograms show disrupted locomotor activity, with activity no longer consolidated to the dark phase in dams exposed to CRD (Fig.1B). While total sleep time is unaffected (*t*<1), sleep is also no longer consolidated to the light phase, as shown by an increase in sleep time in the dark, or active phase (*t*_(9)_=3.614, *p*=0.0056), and a decrease in the light, or inactive phase (*t*_(9)_=4.98, *p*=0.0008)(Fig.1C).

**Figure 1.**
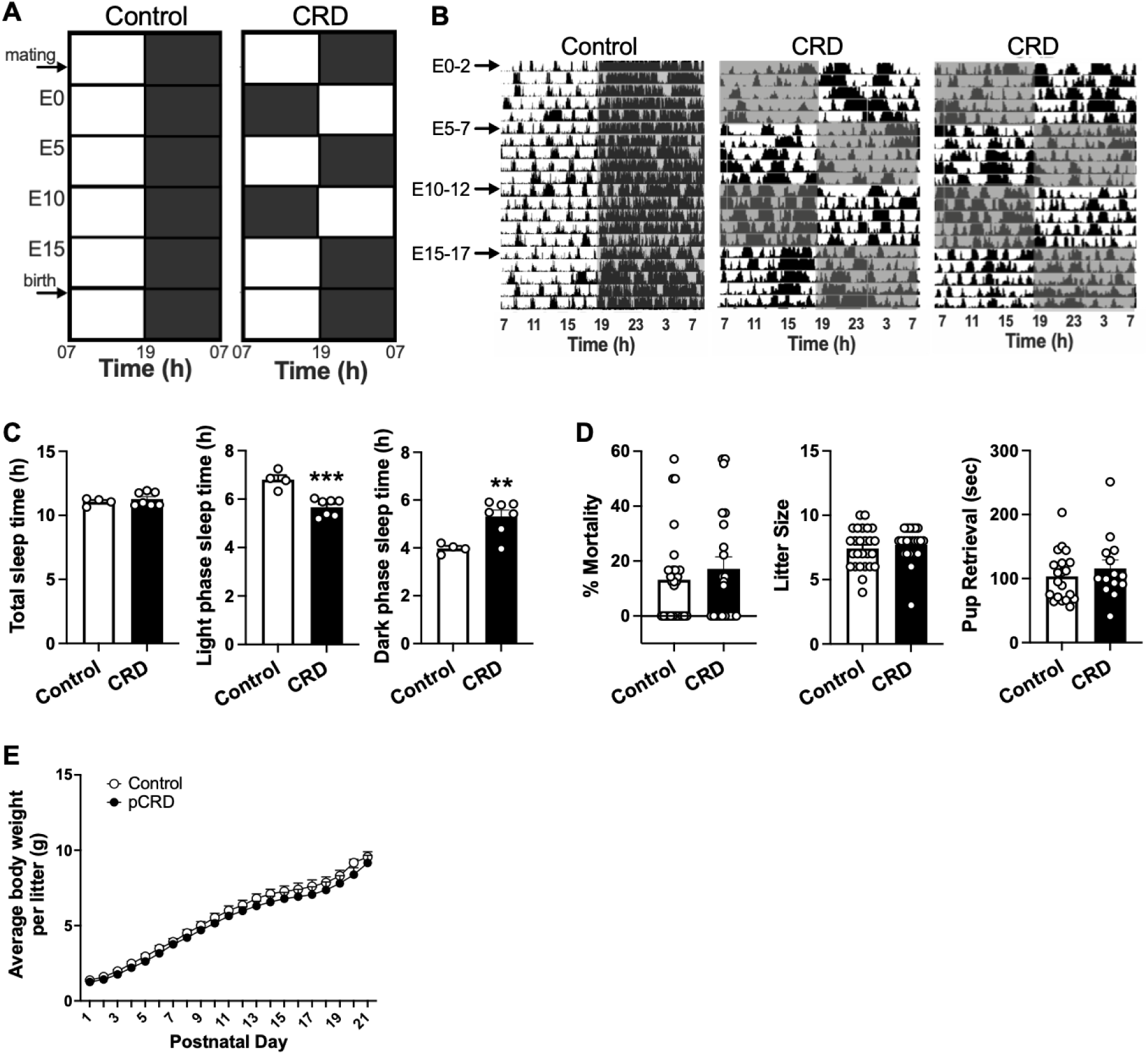
CRD affects locomotor activity and sleep in dams. **(A)** Pregnant dams were exposed to circadian rhythm disruption (CRD) via 4, 12-h light/dark (LD) cycle shifts every 5 days during gestation. **(B)** Locomotor activity rhythms and **(C)** sleep were affected in CRD dams, with activity and sleep no longer consolidated to the appropriate time of day (TOD). **(D)** Pregnancy outcomes, such as % pup mortality and litter size, as well as maternal care, were not affected by CRD. **(E)** Body weight, averaged within litter, was also unchanged after pCRD. Mean+SEM, *n*=4-7 dams (C), 14-26 pups (D), 4-5 litters (E), individual data points are shown on bar graphs, \*\**p*<0.01, \*\*\**p<*0.001.

We also examined pregnancy outcomes, maternal care and body weight to ensure offspring developed normally after CRD. Pup mortality, litter size and pup retrieval were all unaffected by CRD (*t*s<1)(Fig.1D). To account for litter effects, body weights were averaged within each litter. All mice gained weight (session: *F*_(20,140)_=377.6, *p*<0.0001) and pCRD did not affect body weight (interaction: F<1, pCRD: *F*_(1,7)_=1.067, *p*=0.336).

### pCRD has sex-specific effects on cocaine self-administration

To determine whether pCRD induces vulnerability towards SU we used intravenous cocaine self-administration to measure reward-related behaviors during both the light phase and the dark phase. Mice were first trained to self-administer food before jugular catheters were placed. A 3-way ANOVA for the effects of pCRD, sex, and session revealed a trending interaction during the light phase (*F*_(4,440)_=2.382, *p*=0.051). Therefore, male and female mice were analyzed using 2-way ANOVAs. A significant session by pCRD interaction was found (*F*_(4,212)_=3.602, *p*=0.0073), suggesting female mice exposed to pCRD respond less for food, particularly in later sessions. In male mice, no effects of pCRD were found (*F*s<1)(Fig.2A). In the dark phase, we found a significant effect of pCRD (*F*_(1,95)_=10.91, *p*=0.0013) and trending effect of sex (*F*_(1,95)_=3.801, *p*=0.0542), with subsequent analyses revealing a significant effect of pCRD in females (*F*_(1,49)_=8.031, *p*=0.0067) and trending effect in males (*F*_(1,50)_=3.772, *p*=0.0578)(Fig.2B). In both cases, pCRD mice responded more for food than controls.

**Figure 2.**
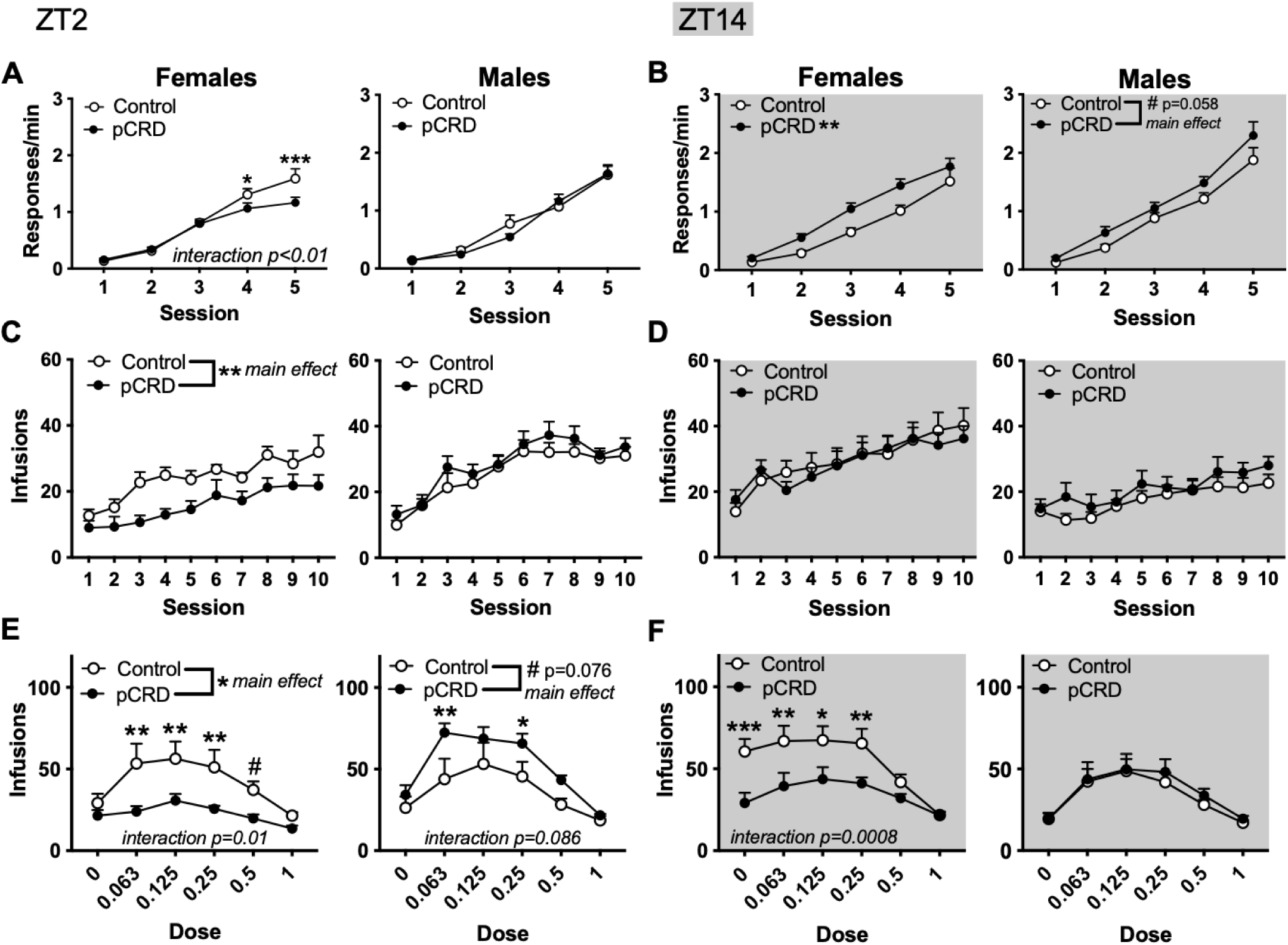
Effects of pCRD on food and cocaine self-administration. **(A)** At ZT2, during the light phase, pCRD reduced lever responding for food in females, but did not affect behavior in males. **(B)** At ZT14, in the dark phase, pCRD increased responding in females, and to a lesser extent, males. **(C)** In females (left), cocaine self-administration was decreased in pCRD mice during light phase acquisition, but no effects were observed in males (right) or in the dark phase **(D)**. **(E)** During a dose response analysis in the light phase, a significant interaction was found where females exposed to pCRD self-administered fewer infusions of cocaine across several doses (left). (Right) In males, a trending interaction and main effect of pCRD were found, suggesting pCRD increased self-administration of cocaine. **(F)** In the dark phase, a similar interaction was found in females with decreased self-administration of cocaine in pCRD mice, particularly at low doses in a dose response curve (left). No effects were found in males (right). Mean+SEM, *n*=21-22 (A), 25-26 (B), 8-12 (C,E), 13-17 (D,F), #*p*<0.1, *\*p*<0.05, \*\**p*<0.01, \*\*\**p<*0.001.

After mice recovered from jugular catheterization, they were trained to self-administer cocaine on the opposite lever. All mice trained in the light phase acquired cocaine self-administration with an increase in lever pressing across sessions (session: *F*_(9,360)_=35.93, *p*<0.0001). A significant effect of sex (*F*_(1,40)_=10.25, *p*=0.0027) and a sex by pCRD interaction (*F*_(1,40)_=6.599, *p*=0.014) were found. When males and females were analyzed separately, we found no effects of pCRD in male mice (*F*s<1), but female pCRD mice self-administered fewer infusions than controls (pCRD: *F*_(1,19)_=8.761, *p*=0.008)(Fig.2C). In the dark phase, there were significant effects of session (*F*_(9,504)_=25.453, *p*<0.0001) and sex (*F*_(1,56)_=11.51, *p*=0.0013), but no effects of pCRD (*F*s<1)(Fig.2D).

To further investigate effects on acquisition, we quantified total drug intake (infusions) and found a pCRD by sex interaction in the light phase (*F*_(1,39)_=4.505, *p*=0.0402)(Supp.Fig1A). Although no significant post-hocs were found, this interaction indicates that male and female mice are oppositely affected by pCRD. In the dark phase, we found an effect of sex (*F*_(1,51)_=11.95, *p*=0.0011), but no effects of pCRD (*F*s<1).

Following acquisition, a dose-response curve was used to measure the reinforcing properties of cocaine. During the light phase, we found a 3-way interaction (pCRD x sex x dose: *F*_(5,160)_=4.507, *p*=0.0007). We then found a 2-way interaction in females (*F*_(5,70)_=3.256, *p*=0.0106), as well as a main effect of pCRD (*F*_(1,14)_=6.112, *p*=0.0269), suggesting reinforcement is decreased in females exposed to pCRD (Fig.2E). In male mice, we found a trending interaction (*F*_(5,90)_=2.002, *p*=0.0859) and effect of pCRD (*F*_(1,18)_=3.542, *p*=0.0761), indicating the opposite or increased reinforcement after pCRD. In the dark phase, a trending effect of sex (*F*_(1,55)_=3.523, *p*=0.0658) and pCRD by dose by sex interaction (*F*_(5,275)_=2.157, *p*=0.0591) were found (Fig.2F). These results are likely driven by female mice, since we found a main effect of pCRD (*F*_(1,27)_=6.346, *p*=0.018) and interaction (*F*_(5,135)_=4.512, *p*=0.0008) in females, but no significant effects were found in males (*F*s<1).

Next, we assessed motivation using a progressive ratio schedule. During the light phase, a 3-way ANOVA revealed a trending interaction (sex x pCRD x dose: *F*_(2,36)_=2.708, *p*=0.0803) and main effect of sex (*F*_(1,18)_=3.124, *p*=0.0941), but no effects of pCRD were found in either sex (female: pCRD *F*_(1,18)_□=□1.792, *p*□=□0.1974, interaction *F*<1; male: pCRD *F*_(1,36)_=1.286, *p*=0.2642, interaction *F*_(2,36)_=1.042, *p*=0.3630)(Fig.3A). In the dark phase, we found a trending interaction (*F*_(2,76)_=2.421, *p*=0.0956) and significant main effect of pCRD (*F*_(1,38)_=7.744, *p*=0.0083) for breakpoint ratio. In females, a significant effect of pCRD was observed (*F*_(1,20)_=5.954, *p*=0.0241) with a decrease in breakpoint ratio, or motivation, in mice exposed to pCRD (Fig.3B). In males, we found a trending interaction (dose x pCRD: *F*_(2,36)_=3.095, *p*=0.0575), but pCRD male mice only responded less at one dose.

**Figure 3.**
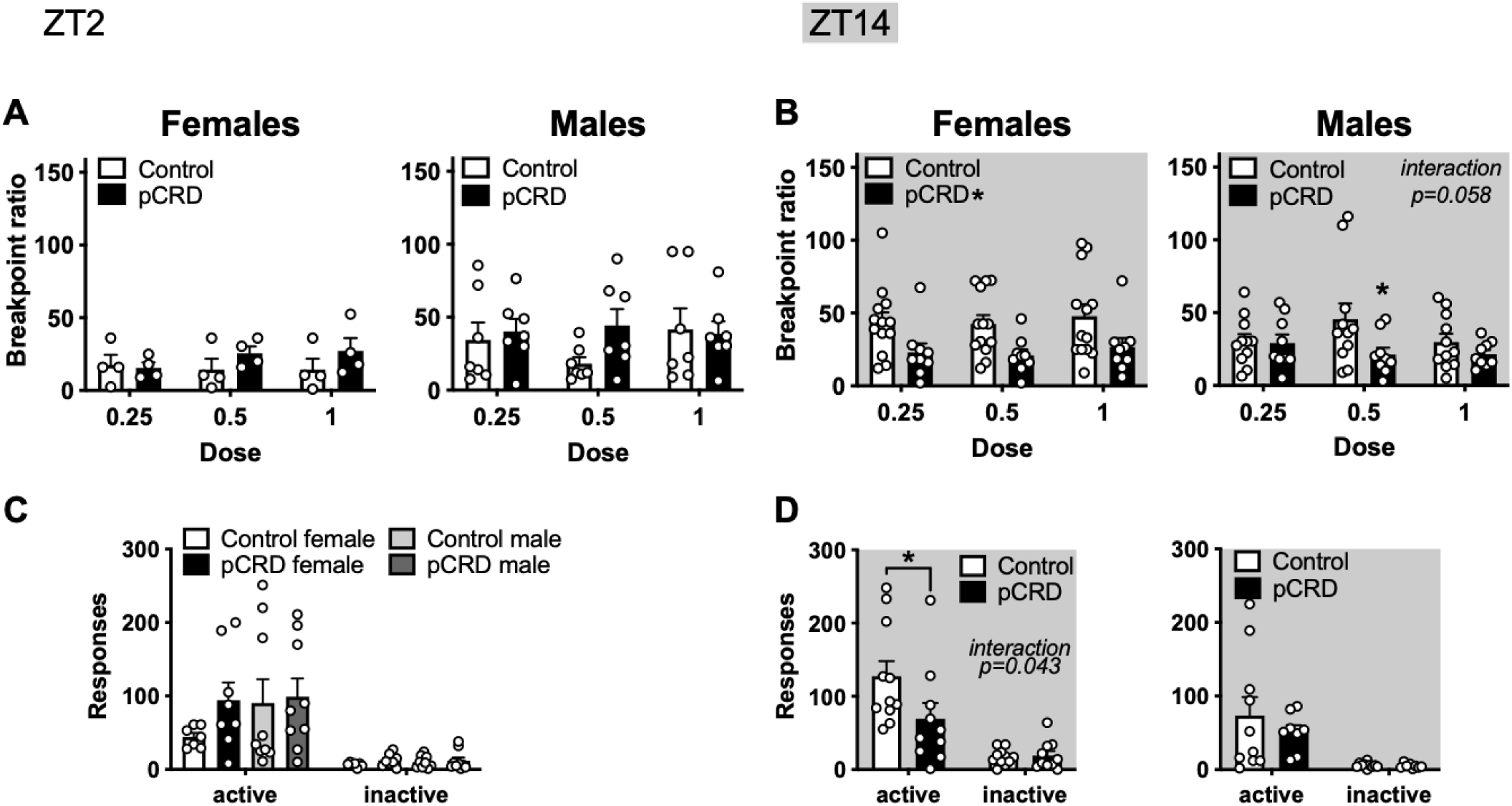
Effects of pCRD on motivation and cue-induced reinstatement. **(A)** There was no effect of pCRD on motivation (breakpoint ratio) to self-administer cocaine during the light phase. **(B)** During the dark phase, motivation was decreased in females with pCRD (left). In males (right), there was a trending interaction between pCRD and dose, with a decrease in motivation found after pCRD, but only for the 0.5 mg/kg/infusion dose of cocaine. For cue-induced reinstatement, **(C)** there was no effect of pCRD during the light phase. **(D)** During the dark phase, a significant interaction was found in females (left), where pCRD mice showed less reinstatement on the previously active lever, but no effect was found in males (right). Mean+SEM, *n*=4-13, *\*p*<0.05.

Mice then underwent extinction testing. Similar effects were found during the light and dark phase, with all mice extinguishing responding (effect of session, light: *F*_(9,270)_=18.05, *p*<0.0001, dark: *F*_(9,342_=35.62, *p*<0.0001). We also found a sex by session interaction in the light phase (*F*_(9,270)_=2.356, *p*=0.0142) and trending sex effect in the dark phase (*F*_(1,38)_=3.622, *p*=0.0646). When males and females were analyzed separately, no effects of pCRD were found during the light (Supp.Fig.1B)(pCRD: *F*s<1, pCRD by session: female *F*_(9,126)_=1.477, *p*=0.1632, male *F*_(9,144)_=1.129, *p*=0.3460) or dark phase (Supp.Fig.1C)(female: pCRD *F*_(1,20)_=2.746, *p*=0.1131, pCRD by session *F*_(9,126)_=1.477, *p*=0.1632; male *F*s<1).

To model relapse-like drug seeking in the absence of a reinforcer, mice underwent cue-induced reinstatement. When previously cocaine-associated cues were re-introduced in the absence of cocaine, all mice demonstrated cue-induced reinstatement by pressing the previously active lever more than the inactive lever in the light phase (lever *F*_(1,29)_=34.69, *p*<0.0001)(Fig.3C). No effects of sex or pCRD were found (*F*s<1). During the dark phase, a 3-way ANOVA revealed a significant effect of lever (*F*_(1,35_=49.34, *p*<0.0001) and sex (*F*_(1,35)_=4.270, *p=*0.0463), as well as a trending effect of pCRD (*F*_(1,35)_=3.490, *p*=0.0701). The effect of pCRD was driven by female mice as no effects were identified in males (*F*s<1), but a session by pCRD interaction in females (*F*_(1,19)_=4.710, *p*=0.0429), indicating reduced cue-induced reinstatement in females exposed to pCRD (Fig.3D). In summary, we found that pCRD decreased cocaine self-administration, motivation across dose and cue-induced reinstatement in females, but increased cocaine intake in a dose response analysis in males.

### pCRD alters reward-related behavior in female mice

We next investigated whether pCRD affects other reward-related behaviors. Here, one sample t-tests were performed to assess preference. In sucrose consumption, only female control mice showed a significant preference relative to 50% (no preference)(*t*_(1,13)_=4.308, *p*=0.0009). No preference was found in males (*t*s<1) or female pCRD mice (*t*_(1,21)_=1.595, *p*=0.1257)(SuppFig.2A). In cocaine CPP, all male mice and female control mice showed a preference for cocaine relative to 0, while female pCRD mice did not (male control *t*_(1,15)_=3.514, *p*=0.0031, female control *t*_(1,15)_=4.923, *p*=0.0002, male pCRD *t*_(1,17)_=3.988, *p*=0.001, female pCRD *t*_(1,12)_=1.625, *p*=0.1301)(SuppFig.2B).

### pCRD enhances food self-administration in male mice

Since a sex difference was found in food self-administration prior to light phase cocaine self-administration, a separate cohort of mice were trained to self-administer food at varying schedules of reinforcement during the light phase. Mice were trained to respond on a lever for a food reinforcer (FR1). A 3-way ANOVA showed all mice acquired the response, increasing lever pressing across sessions (F_(4,112)_=51.46, *p*<0.0001)(Fig.3A). A trending interaction (session x pCRD x sex F_(4,112)_=2.09, *p*=0.087) and main effect of sex (F_(1,28)_=3.27, *p*=0.081) were also found. Therefore, 2-way ANOVAs were performed for males and females separately. In male mice, a significant main effect (pCRD F_(1,13)_=5.57, *p*=0.035) and interaction (pCRD x session F_(4,52)_=3.26, *p*=0.019) were observed, with an increase in lever pressing in male pCRD mice late in acquisition. No changes were found in female mice (pCRD *F*<1, interaction F_(4,60)_=2.02, *p*=0.104).

On an FR3 schedule (SuppFig.2C), a significant effect of sex (F_(1,28)_=27.24, *p*<0.0001) was found, but no effects of pCRD were observed in any group (*F*s<1). We then assessed reversal learning and a 4-way ANOVA found that all mice learned the new lever association (lever x session F_(2,56)_=198.67, *p*<0.001). There was also a lever by sex interaction (F_(1,28)_=8.94, *p*=0.006). Subsequent analyses found no effect of pCRD in female mice, while males had a trending lever by pCRD interaction (F_(1,26)_=3.42, *p*=0.076)(Fig.3B). This may indicate that active and inactive lever pressing is differentially affected in control and pCRD mice, but only in males.

Next, mice were trained to respond on an RI schedule and a session by sex by pCRD interaction was found (F_(4,112)_=2.795, *p*=0.0295). When analyzed separately, no effects were found in female mice (*F*s<1), but a trending session by pCRD interaction (F_(4,52)_=2.423, *p*=0.0598) and significant effect of pCRD (F_(1,13)_=9.137, *p*=0.0098) were found in male mice, with increased response rates in male pCRD mice, especially during RI60 (Fig.4C;Supp.Fig2D).

**Figure 4.**
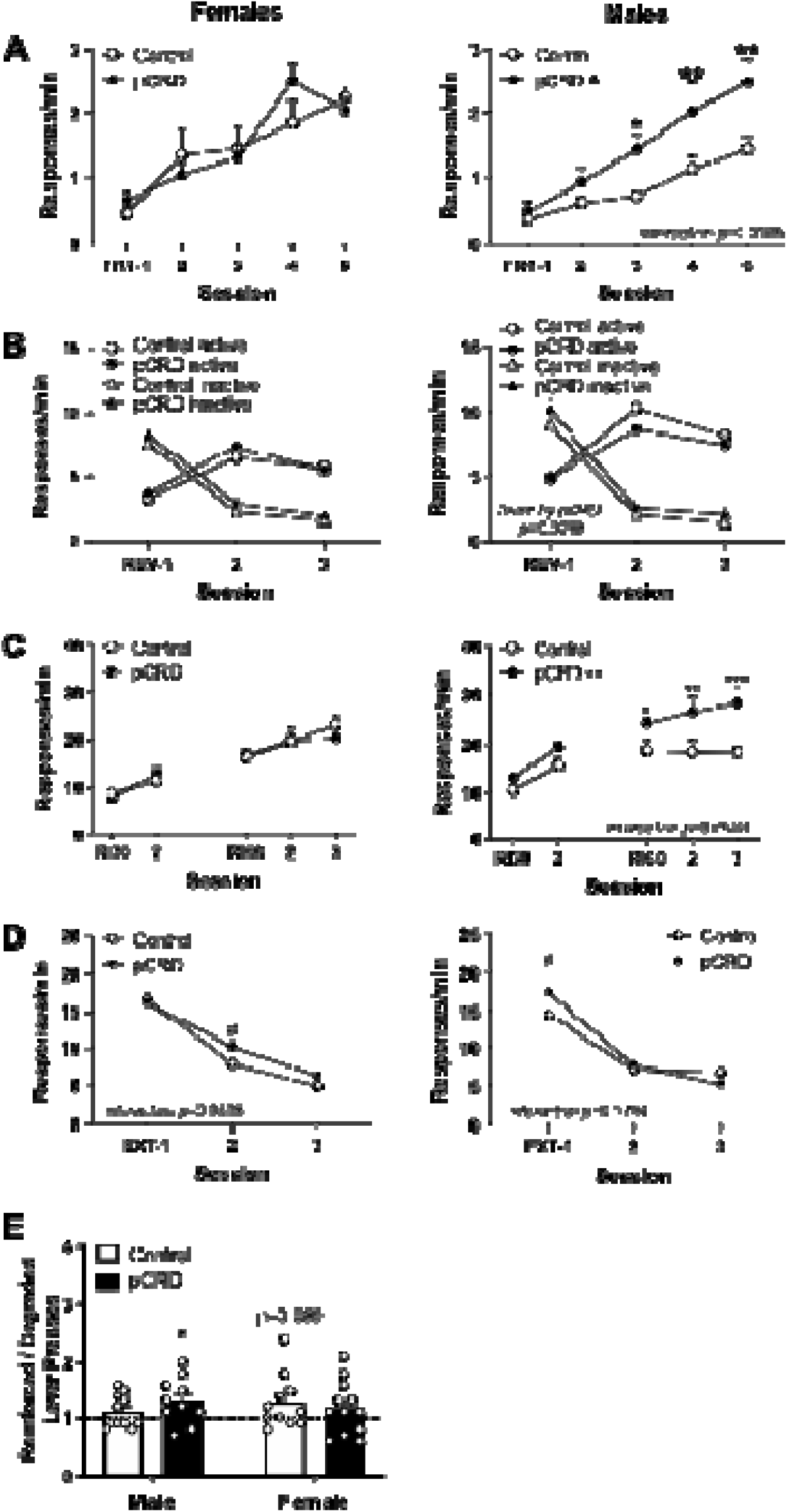
pCRD increases goal-directed action selection in male mice. Operant food self-administration was assessed with a variety of schedules of reinforcement. **(A)** During a fixed ratio 1 (FR1) schedule, operant responding for food reinforcers was unchanged in female mice (left), while male pCRD mice showed higher response rates, especially later in training. **(B)** No effects of pCRD were found in females mice during reversal learning (left), but a trending lever by pCRD interaction was found in males. No significant post-hocs were observed, but this interaction suggests that responding on the previously active and currently active levers is differentially affected by pCRD. **(C)** During random interval (RI) training, again no effects were identified in females, but a significant main effect and interaction were found in males. Specifically, male pCRD mice responded more for food during RI60 training. **(D)** Under extinction conditions, a trending interaction of pCRD by session was found in both females and males, with a trending increase in pCRD mice on only one day of extinction in both males and females. **(E)** A separate cohort of mice were trained on action outcome contingency degradation. While male control mice showed habitual response strategies (lever press ratio of ∼1), pCRD male mice maintained goal-directed response strategies (ratio > 1). In female mice, controls had a trending ratio of greater than 1, suggesting a trend towards goal-directed responding, while pCRD exposed females were habitual. Mean+SEM, *n*=7-9 (A-D), 10-12 (E), #*p*<0.1, *\*p*<0.05, \*\**p*<0.01, \*\*\**p<*0.001.

Under extinction conditions, a session by sex by pCRD interaction was found (F_(2,56)_=4.902, *p*=0.011). We then found trending session by pCRD interactions in males (F_(2,26)_=2.797, *p*=0.0794) and females (F_(2,30)_=3.021, *p*=0.0638)(Fig.3D;Supp.Fig2D). Female mice exposed to pCRD responded more than control mice only on day 2 of extinction, while male pCRD mice responded more on day one of extinction.

In a separate cohort of mice, decision making strategies were investigated using action-outcome contingency degradation, wherein the contingency between one action and its outcome is degraded, while one is maintained (reinforced). This relationship is then probed and mice that equally engage both levers (ratio of 1), regardless of reinforcement status, are considered habitual, while mice that preferentially engage the reinforced lever are considered goal-directed in their response strategy. We found that our training protocol was sufficient to induce habitual response strategies in male control mice (*t*_(1,10)_=1.213, *p*=0.253), while male pCRD mice are goal-directed (*t*_(1,9)_=2.348, *p*=0.044). In female mice, controls tend towards goal-directed decision-making (*t*_(1,10)_=1.885, *p*=0.089), while pCRD mice are habitual (*t*_(1,11)_=1.271, *p*=0.230)(Fig.3E).

### pCRD alters anxiety- and depressive-related behavior sex-specifically

We next investigated whether pCRD affects anxiety- and depression-related behaviors. We first determined whether pCRD affected locomotor response to novelty (Fig.4A). A 2-way ANOVA revealed a significant effect of sex, with higher locomotion in females (F_(1,95)_=5.831, *p*=0.0178), but no effects of pCRD were found (pCRD Fs<1, interaction F_(1,95)_=1.737, *p*=0.1907). While only a trending effect of sex was found in the OFT (F_(1,88)_=3.453, *p*=0.0665, pCRD and interaction Fs<1)(Fig.4B), a significant sex by pCRD interaction was found in the EPM (F_(1,88)_=6.203, *p*=0.0146)(Fig.4C) and LD box (F_(1,95)_=4.295, *p*=0.0409)(Fig.4D). Specifically, male pCRD mice spent more time in the lit compartment of the LD box, indicating increased risk taking or exploratory drive. In contrast, females showed decreased open arm entries in the EPM, indicating increased anxiety-like behavior in female pCRD mice. We next assessed the effect of pCRD on helplessness-like behavior using the FST and found a significant interaction (F_(1,60)_=6.371, *p*=0.0143)(Fig.4E). Post-hoc analyses revealed that male pCRD mice had reduced immobility compared to controls, suggesting pCRD may reduce depression-related behavior in males only.

### Cross fostering does not alter pCRD-induced behavioral phenotypes

Cross fostering was used to investigate whether altered maternal care might be driving the behavioral phenotypes identified above. We investigated effects on a subset of behaviors and, overall, found similar patterns suggesting that maternal care in CRD dams is not affecting behavioral outcomes in pCRD mice. In CPP, all groups had a significant preference for cocaine using one-way t-tests (female control *t*_(1,15)_=6.082, *p*<0.0001, female pCRD *t*_(1,15)_=2.777, *p*=0.0141, male control *t*_(1,19)_=8.030, *p*<0.0001, male pCRD *t*_(1,19)_=10.29, *p*<0.0001), therefore, we performed a 2-way ANOVA and found a sex by pCRD interaction (F_(1,68)_=11.78, *p*=0.0010)(Fig.4F). Post-hocs indicated a decreased preference for cocaine in female pCRD mice and an increased preference in males. For anxiety-related behavior, we found a trending sex by pCRD interaction in the OFT (F_(1,83)_=3.30, *p*=0.0729)(Fig.4G). Although no significant post-hocs were found, this interaction indicates that males and females are differently affected by pCRD after cross fostering. No effects were found in EPM (*F*s<1, interaction F_(1,85)_=1.241, *p*=0.2683)(Fig.4H), but the direction of effect is consistent with prior findings (Fig.5C).

**Figure 5.**
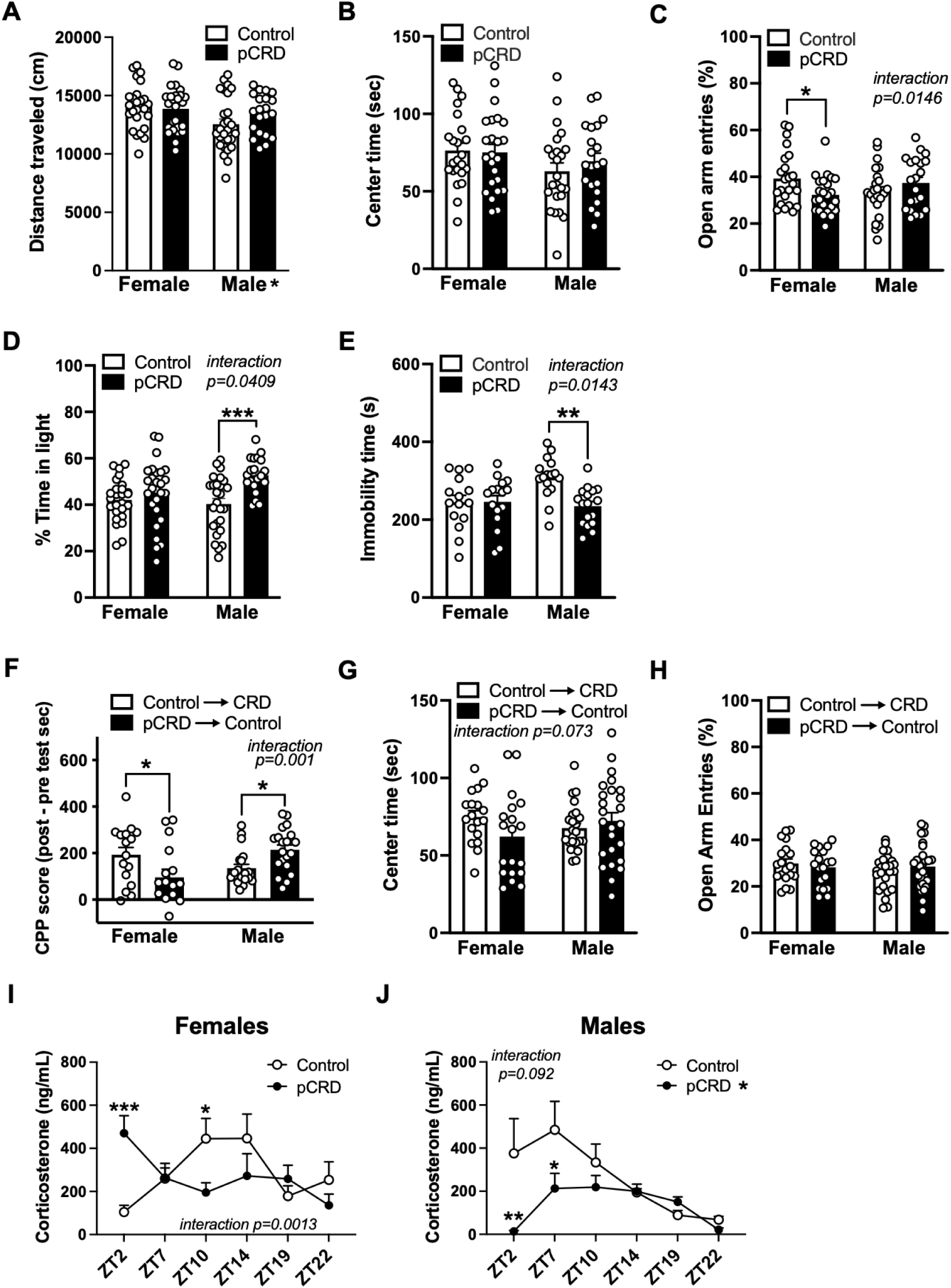
pCRD sex-specifically affects anxiety- and depressive-related behavior and corticosterone rhythms. **(A)** pCRD did not affect the locomotor response to novelty, but a sex difference was found where females were more active than males. **(B)** While no effects of pCRD were found in the open field test (OFT), **(C)** a significant interaction was found in the elevated plus maze (EPM) with decreased open arm entries in pCRD females. **(D)** Similarly, an interaction was found in the light/dark box **(E)** and forced swim test, but here a decrease was observed in anxiety-related behaviors, as shown by an increase in light time, and helplessness behavior, in male pCRD mice. **(F-H)** The effects of cross-fostering on pCRD-induced behavioral changes were investigated. Control➔pCRD = pups born to control dams and cross-fostered to pCRD dams at birth. pCRD➔Control = pCRD born pups raised by a control dam. **(F)** After cross-fostering, we saw similar effects of pCRD on cocaine conditioned place preference (CPP), with a significant interaction and post-hoc tests indicating that CPP is reduced in pCRD females, but increased in pCRD males. **(G)** A similar effect was found in the OFT, with a trending sex by pCRD interaction. No post-hoc effects were found, but this interaction suggests that center time is differentially regulated by pCRD in males and females; females tend to spend less time in the center and males more time after pCRD. **(H)** No significant effects of pCRD were found in the EPM after cross-fostering. Rhythms in corticosterone were measured across 6 times of day. **(I)** In females an interaction revealed that while rhythms in corticosterone typically peak around ZT10 in control mice, after pCRD rhythms shift with a decrease in concentration at ZT10 and an increase at ZT2. **(J)** In males, an interaction was also found that indicates a dampening of rhythms in pCRD mice, with lower concentrations early in the light phase in pCRD males compared to controls. Mean+SEM, *n*=20-27 (A-D), 15-17 (E), 16-26 (F-H), 5-8 (I,J), *\*p*<0.05, \*\**p*<0.01, \*\*\**p<*0.001.

### pCRD affects rhythms in corticosterone sex-dependently

To investigate whether pCRD might be mediating behavioral changes through mechanisms involving the stress hormone corticosterone (CORT), we measured rhythms in CORT as well as the glucocorticoid response to acute stress. We first confirmed that CORT rhythms were present in both male and female control mice. We found an effect of time (F_(5,74)_=2.798, *p*=0.0227) indicating that CORT levels vary across TOD, as well as a time by sex interaction (F_(5,74)_=3.034, *p*=0.0151)(Supp.Fig.3A). Post-hocs showed that CORT rhythms vary by sex, where females have a slightly later peak time, around dusk (ZT10-14), compared to earlier in the light phase (∼ZT7) in males. However, this small phase shift across sex could be due to slightly different waveforms or amplitudes, as well as more variability in male control mice.

A 3-way ANOVA revealed a sex by pCRD by TOD interaction (F_(5,136)_=4.899, *p*=0.0004) (Supp.Fig.3B). Data were separated by sex and 2-way ANOVAs were performed. In females, a significant TOD by pCRD interaction was found (F_(5,72)_=4.469, *p*=0.0013) with higher CORT concentrations at ZT2 and lower at ZT10 in pCRD mice (Fig.4I). In males, a trending interaction was found (F_(5,64)_=1.987, *p*=0.0924), as well as a main effect of pCRD (F_(1,64)_=6.289, *p*=0.0147) where CORT concentrations were lower after pCRD, particularly early in the light phase, when concentrations tend to peak in controls (∼ZT7)(Fig.4J).

We next investigated whether the glucocorticoid response to acute stress is altered after pCRD. In a 3-way ANOVA only an effect of time (F_(3,120)_=8.063, *p*<0.0001) was found, indicating that CORT levels responded to acute stress (Supp.Fig.3C). No significant effects of sex or pCRD were found, indicating the acute stress-induced change in CORT did not differ after pCRD.

## Discussion

Here, we aimed to investigate the effects of pCRD on reward- and mood-related behavior to better understand how disrupted rhythms during gestation affect behavior in adulthood. We found that pCRD produces long-lasting, sex-dependent changes in reward- and affective-like behaviors, as well as corticosterone rhythms.

Overall, we found sex-specific phenotypes after pCRD where males exhibited increased reward sensitivity and risk-taking behavior, while females had decreased reward sensitivity and increased anxiety-like behavior. We used a myriad of reward- and affective-related tasks to investigate phenotypic changes. Females exposed to pCRD not only self-administered fewer infusions of cocaine during acquisition in the light phase, but also in a dose-response analysis in the light and dark phase. During the dark phase, pCRD females also had a substantial decrease in motivation, as shown by a reduced breakpoint ratio for cocaine, and in cue-induced reinstatement, which is a model for cue-induced relapse. In contrast, males self-administered more infusions of cocaine during a dose-response analysis in the light phase. These findings suggest that females have a profound decrease in reward sensitivity, while males have a mild increase, which is more sensitive to time of day (TOD).

We next examined whether these findings extend to other reward-related behaviors. Reward preference was only affected by pCRD in female mice; while control females had a significant preference for sucrose and cocaine, pCRD mice showed no preference. Food self-administration, however, was more affected in males, with increased responding during an FR1 and RI60 schedule of reinforcement in male pCRD mice. We also found a trending interaction during reversal learning, suggesting lever responding was affected differently in control and pCRD males, indicating increased perseveration on the previously active lever, as well as impaired cognitive flexibility (decreased responding on the new active lever) in male pCRD mice. Male pCRD mice also maintained goal-directed decision making after extended training that elicits habitual response strategies in control mice. All behavioral responses found in male pCRD mice (*e.g.* increased reward sensitivity, perseveration) could be driven by stronger, sustained goal-directed decision making, indicating male pCRD mice might be particularly driven by outcomes.

In female pCRD mice, there was an overall decrease in reward sensitivity across modality, but responding to food reinforcers in operant conditioning was largely unaffected. To determine whether decreased reward sensitivity reflects a resiliency towards SU, or alternatively a broader phenotypic change, such as an anhedonic-like phenotype, we extended testing to affective-like behaviors. We found that pCRD males with increased reward sensitivity also had increased exploratory drive, suggesting increased risk taking, as well as more active coping in the FST. Together these findings further demonstrate that males exposed to pCRD might be vulnerable to SU and associated behaviors. In contrast, pCRD females with decreased reward sensitivity also had increased anxiety-like behavior. Considering reduced cocaine intake, motivation, cue-induced reinstatement, cocaine preference, sucrose consumption and increased anxiety-like behavior, pCRD females might be vulnerable to anhedonia and associated behavioral changes.

Although we found no effects of our pCRD model on maternal care and early postnatal development, previous studies have identified these changes. For example, a paradigm with 6-h shifts every 4 days from E0-P20^45^ alters pup retrieval and anxiety-related behavior. In this case, shifted pups cross-fostered to shifted dams were most affected, followed by shifted pups cross-fostered to control dams and then control pups cross-fostered to shifted dams, suggesting the gestational effects of CRD contribute more to behavioral outcomes, but postnatal effects also play a role. However, in this model, disrupted mice were actively exposed to CRD during early postnatal development making it difficult to determine whether maternal care, or behavior in control mice cross-fostered to shifted dams, are affected because of gestational CRD or ongoing CRD. Here, we attempted to limit postnatal effects by isolating CRD to gestation. We found persistent behavioral effects after pCRD despite cross-fostering, further indicating that the prenatal environment, including *in utero* hormones or circadian cues, are likely mediating effects of pCRD in our model.

Sexual dimorphism observed after pCRD could be regulated by a variety of factors, including chromosomal sex, gonadal hormones (both organizational hormnes during development and activational hormones in adulthood) and underlying circadian differences. Developmental hormones could contribute to the effects of pCRD since exposure to gonadal hormones during critical periods induce long-term changes to the brain and behavior. For instance, prenatal androgen activation sex-specifically alters adult alcohol consumption and nucleus accumbens gene expression^53^. We therefore hypothesize that pCRD interacts with developmental hormones *in utero* to alter reward sensitivity sex-dependently. In future studies, we will use the four core genotypes mice^54^ to differentiate effects of genetic and gonadal sex and, coupled with gonadectomy, determine whether developmental or adult hormones are driving behavior.

The timing of gonadal hormone exposure during development may be especially important for these sex differences. In males, a surge in androgens, produced by the developing testes, occurs during late gestation (∼E16-18) and is critical for sexual differentiation in the brain across species^55,56^. Notably, this timing closely aligns with the final LD cycle shift in our model, when rhythms are highly disrupted in CRD dams. Therefore, disruption of rhythms during this period may interfere with the timing or magnitude of this developmental surge. Future studies are needed to determine whether pCRD-induced behavioral alterations depend on disruption during early versus late gestation.

In contrast, females do not experience a comparable surge, suggesting that pCRD effects may arise through distinct mechanisms. Placental function is sexually dimorphic, with differences in gene expression, hormone regulation and signaling, as well as responses to maternal stress, resulting in female placentas generally exhibiting greater adaptive capacity than males^57,58^. Alternatively, pCRD-induced behavioral changes might be caused by long-term changes in circadian rhythms, which are controlled by the central pacemaker or suprachiasmatic nucleus (SCN). It is well known that sex differences exist in circadian rhythms, within the SCN^59^, as well as in the extra-SCN brain in humans^60^ and rodents^61^. We expect that while androgen-dependent mechanisms might drive effects of pCRD in males, sexually dimorphic placental signaling and/or long-term reorganization of circadian systems drive effects in females.

pCRD may reorganize the circadian system through altered maternal entrainment of the fetal SCN, as maternal signals are known to drive rhythms *in utero*^62^. A recent study confirmed that daily rhythms are detectable *in utero*, developing gradually and stabilizing by ∼E15.5^63^, highlighting a sensitive window during which maternal disruption may have lasting effects. One mechanism that may mediate the effects of pCRD in both sexes is glucocorticoid signaling. Maternal glucocorticoids are implicated in fetal development, and recent work demonstrates that exposure to CORT during late gestation phase shifts fetoplacental rhythms *in utero*, while blocking signaling reduces synchrony between maternal and fetal placenta^63^. Thus, glucocorticoids may play a role in how rhythms develop and synchronize with the mother prior to birth. While we did not observe effects of pCRD on the glucocorticoid response to acute stress, indicating that overall HPA axis reactivity is largely intact, we did identify sex-specific changes in CORT rhythms. In females, pCRD shifted rhythms in CORT, changing peak times from ∼ZT10 to ZT2. In males, pCRD reduced CORT levels, dampening rhythms by decreasing concentrations early in the light phase when levels would typically peak. These distinct alterations may contribute to the sex-specific behavioral phenotypes observed after pCRD. Prior studies have found blunted CORT rhythms following pCRD, however, these measurements were obtained prenatally, a more severe CRD paradigm was used, and sex differences were not assessed^64^. When both sexes were examined following exposure to this greater level of pCRD, diurnal rhythms in CORT were lost in both sexes in adulthood. However, CORT was only measured at two TOD, ∼ZT2-4 and ∼ZT14-16, which may obscure phase shifts^65^. Our findings suggest pCRD may differentially alter glucocorticoid rhythmicity in males and females, with blunted rhythms in males and shifted rhythms in females. Together, these data support a model in which pCRD disrupts maternal-fetal circadian alignment, at least in part through glucocorticoid-dependent mechanisms, with downstream consequences that diverge by sex.

Overall, these experiments found that LD cycle disruptions during gestation altered reward-related behavior sex-specifically. Female mice exposed to pCRD showed decreased cocaine intake, motivation, cue-induced reinstatement, cocaine and sucrose preference, and increased anxiety-related behaviors. In contrast, male pCRD mice showed increased cocaine intake, food self-administration, including perseveration and sustained goal-directed decisioning making, and exploratory drive/risk taking. We also found sex-specific effects on corticosterone rhythms, where female pCRD mice have a shift in peak CORT levels, while male pCRD mice have dampened rhythms. These findings demonstrate that CRD during pregnancy produces long-lasting, sex-dependent alterations in reward- and mood-related behavior, potentially mediated by changes in circadian and endocrine signaling.

## Acknowledgements

The author(s) declare that financial support was received for the research, authorship, and/or publication of this article. This work was funded by the National Institutes of Health: DA039865, DA046346, MH106460, MH111601 (PI: Colleen McClung) and DA046117, DA055064, L60DA054665 (PI: Lauren DePoy). Additional funding was provided by the Brain and Behavioral Research Foundation (29386, DePoy), and the WoodNext Foundation (PI: McClung). Cocaine was provided by NIDA via the NIH drug distribution center.

## Conflicts of interest

The author(s) have no conflicts of interest to disclose.

## Notes

### Competing Interest Statement

The authors have declared no competing interest.

