## Supplemental Figures for "Prenatal circadian rhythm disruption induces sex-specific substance use and mood-related phenotypes in mice"

Supplementary Figures

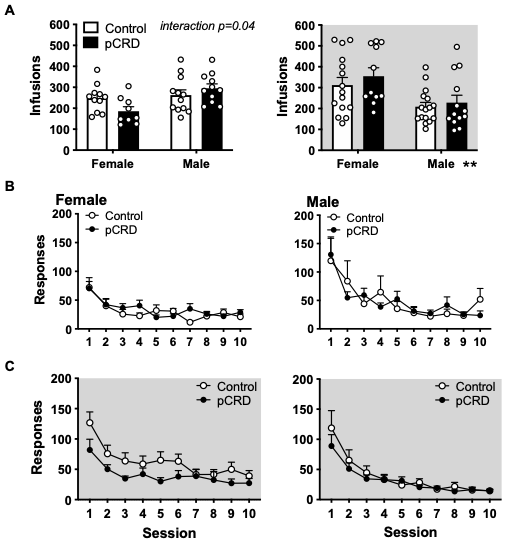

**Supplementary Figure 1. Effects of pCRD on cocaine intake and extinction. (A)** During the light phase, there was a significant pCRD by sex interaction. Although no significant post-hocs were found, this suggests an opposite effect of pCRD on drug intake in males and females (left). During the dark phase, there was no effect of pCRD, but a significant effect of sex indicating that females self-administered more cocaine than males (right). pCRD did not affect responding under extinction conditions in the light phase **(B)** or dark phase **(C)**. Mean+SEM, *n*=9-16 (A), 7-12 (B,C) ***p*<0.01.

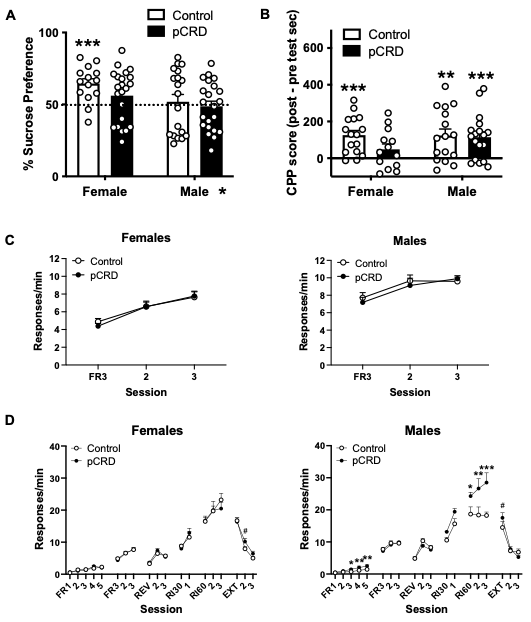

**Supplementary Figure 2. Sex-specific effects of pCRD on reward. (A)** Female pCRD mice have a reduced preference for a sucrose solution, as indicated by a lack of preference for sucrose over water, while female control mice have a significant preference for sucrose. No changes were found in males. **(B)** Female pCRD mice did not develop a preference for cocaine in CPP, while all other groups showed a significant preference (compared to 0). During food self-administration, **(C)** there were no effects of pCRD on fixed ratio 3 (FR3) responding. **(D)** All food training behaviors are shown on the same chart for ease of visualization in females (left) and males (right). Mean+SEM, *n*=14-22 (A), 13-18 (B), 7-9 (C,D), #*p*<0.1, **p*<0.05**,** ***p*<0.01, ****p<*0.001.

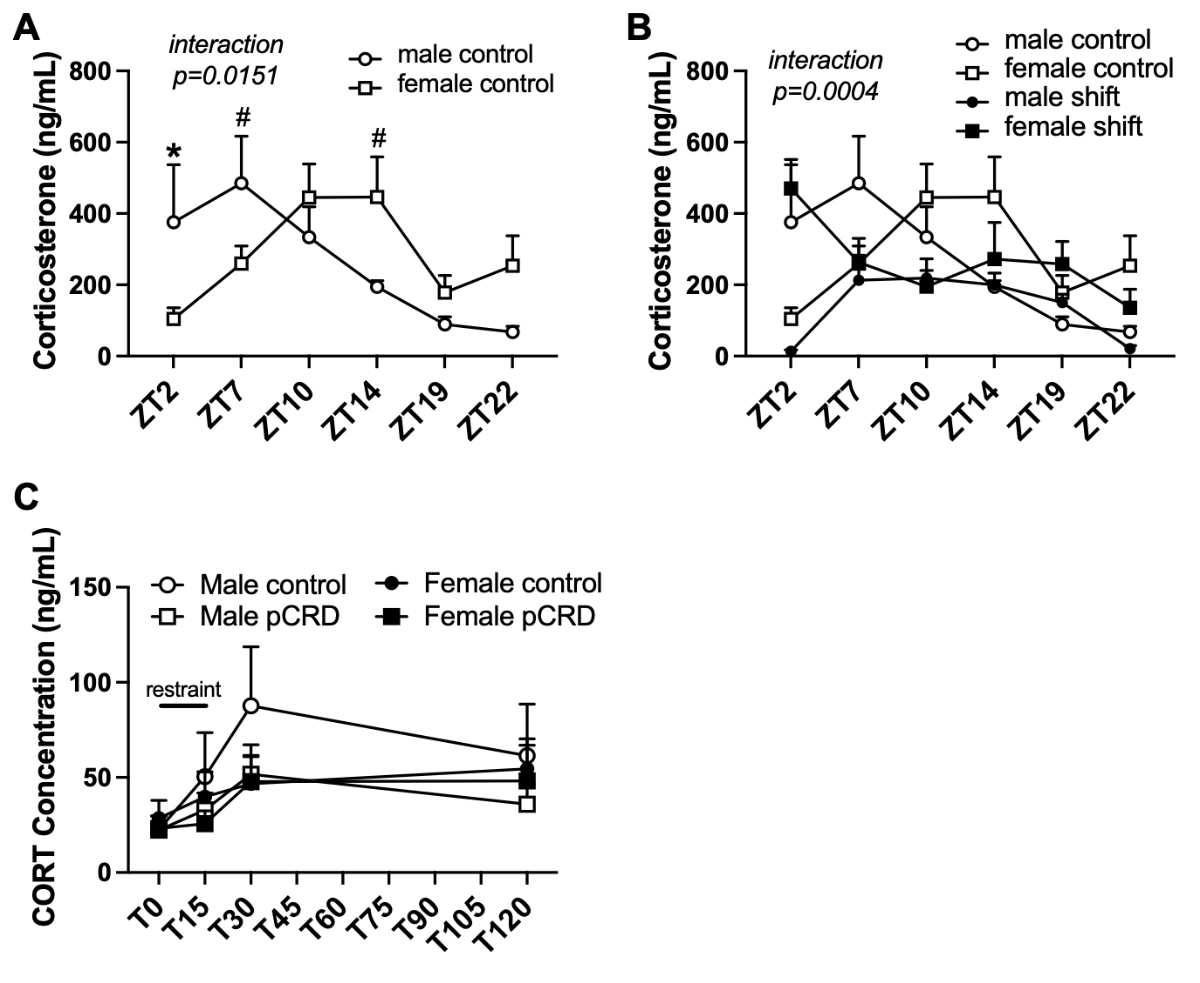

**Supplementary Figure 3. Effects of pCRD on corticosterone rhythms and response to stress. (A)** Control males and females have different levels of corticosterone across time of day. In males, corticosterone concentration peaks early in the light phase, while levels peak later in the day. **(B)** Corticosterone rhythms are shown for males and females with or without pCRD. **(C)** There was no effect of pCRD on the glucocorticoid response to acute restraint stress. Mean+SEM, *n*=5-8 (A,B), 8-13 (C), #*p*<0.1, **p*<0.05.
