## Supplemental Methods for "Prenatal circadian rhythm disruption induces sex-specific substance use and mood-related phenotypes in mice"

Supplementary Methods

**Animals and housing conditions**

Male and female C57BL/6J mice were acquired from The Jackson Laboratory (000664) and subsequently bred in-house. The mice were housed under a 12-hour light/dark (LD) cycle, with lights turning on at 0700 (Zeitgeber Time, ZT0), except during circadian disruption. Behavioral assessments were conducted during the light phase, between ZT2 and ZT7, unless otherwise noted. Food and water were provided ad libitum unless otherwise indicated. All procedures were approved by the University of Pittsburgh or University of Toledo Institutional Animal Care and Use Committee.

**Circadian rhythm disruption**

CRD was initiated in 8-12 weeks old pregnant dams. Three days after harem breeding pairs were established, dams were separated and shifted 12-hour from a normal 12:12 light/dark (LD) cycle to a 12:12 schedule with lights on at 1900. After 5 days, the light cycle was shifted by 12-hour back to a 12:12 schedule with lights on at 0700. The 12-hour shifts occurred every 5 days during gestation, resulting in 4 complete reversals of the LD cycle before birth. Control animals were kept on a normal light schedule, but cages were sham handled. Offspring were born under a normal LD cycle (ZT0 = 0700). Behavioral tests were performed in adult offspring between 8 and 20 weeks of age.

Birth outcomes were tracked, including pup mortality and litter size. Pup retrieval was also measured at ~P7, as a proxy for maternal care. In brief, the dam was removed from the cage, while the pups were spread equidistantly around the cage. Latency to return all pups to the nest was measured. In one cohort, body weight was also measured at each postnatal day (P) until P21. Tattoo ink was used to semi-permanently mark pups. Markings were refreshed every 4-5 days and dams were marked to ensure pups were still accepted by the mother despite any ink odor.

Homecage activity and sleep/wake measurements were recorded for a subset of dams during gestation using piezoelectric recording of movements and breath rate (PiezoSleep, Signal Solutions, LLC, Lexington, KY, United States). Dams were individually housed in four-cage unit polycarbonate boxes in a ventilated, light-controlled tent. Each cage rested on a PVDF square sensor (17.8 x 17.8 cm, 110 µm thick) protected by a plastic tray (50.8 µm)^33,34^ with a rubber pad between each sensor and the base to prevent crosstalk between cages. Sensors were connected to an amplifier and pressure signals and breath rates were classified as movements related to activity and inactivity or sleep and wake.

**Behavioral testing**

Mice used for behavioral testing were assigned to only one of six different cohorts, which are described briefly. All behavior was conducted during the light phase at ~ZT2, except for cohort 3, which were trained during both the light phase (~ZT2) and dark phase (~ZT14). Mice were acclimated to testing rooms for 30 minutes before behavioral testing.

*Cohort 1: Exploratory drive*

Mice underwent a battery of tests in the following order to evaluate locomotor activity and exploratory behavior using open field, elevated plus maze (EPM), and light/dark (LD) box as well as depressive-like behavior (forced swim test, FST). Testing was conducted every other day.

**Elevated plus maze**

Mice were placed at the center of the maze, oriented toward an open arm, under dim lighting (~20 lux). The EPM included 2 open and closed arms (30 x 5 cm) perpendicular to each other and elevated 81 cm above ground. Behavior was recorded for 10 minutes. EthoVision XT was used to assess the time spent in the open arms and the number of entries into each arm.

**Light/dark box**

Transparent enclosures (Kinder Scientific Smart Cage Rack System; field dimensions: 9.5″ x 18.0) were partitioned into two equal chambers: a black, opaque chamber covered with a lid to maintain darkness and a brightly illuminated chamber (~880 lux). A doorway connected the two chambers. Mice were initially placed in the dark chamber for 2 minutes before the door was opened, allowing them to explore both sides freely for 20 minutes. Photobeams recorded the number of entries into, and the duration spent in the illuminated chamber.

**Forced swim test**

A subset of mice were individually placed in a glass beaker filled with water (25°C; 18 cm deep), with visual barriers separating the beakers to prevent mice from seeing one another during testing. Behavior was recorded for 10 minutes. A trained observer, blinded to the experimental conditions, scored the latency to immobility and the duration of struggling during the final 8 minutes. Struggling was defined as any movement not solely aimed at staying afloat. Immobility time was calculated by subtracting the time spent struggling from the total 8-minute observation period.

*Cohort 2:* *Reward-related behaviors*

**Conditioned Place Preference**

As previously described^35^: on day 1, a pre-conditioning test was performed, during which mice were placed in the center of a three-chamber box. The two outer chambers were both visually and physically distinct. Time spent in each chamber was recorded over a 20-minute session. For the following 4 days, mice received injections of either saline or cocaine and were confined to one of the outer chambers. Saline was administered on days 2 and 4, while cocaine was administered on days 3 and 5 (5 mg/kg, 10 mL/kg, NIDA drug consortium). A biased design was implemented, as more than 50% of the mice exhibited a chamber preference during the pre-test. The preferred chamber (>10% preference) was paired with saline, while chamber assignments were determined pseudo-randomly for mice that showed no side preference.

**Sucrose preference**

Mice were acclimated to two bottles containing either water or a sucrose solution (0.5% sucrose w/v). After 24 hours, the bottle positions were switched to reduce side bias. 24 hours later, mice were individually housed and given access to both water and sucrose bottles. They remained undisturbed overnight for 18 hours, after which the bottles were weighed to measure sucrose and water consumption. To control for differences in bottle weight due to leakage, control cages underwent sham handling, and fluid loss was recorded. The percentage of sucrose preference was calculated, with 50% indicating no preference.

*Cohort 3:* *Cocaine self-administration*

**Intravenous cocaine self-administration**

Mice were first trained to self-administer food pellets on an FR1 schedule. One cohort was trained in the light phase (~ZT2) and another in the dark phase (~ZT14). Sessions lasted 60 minutes or until 30 pellets were earned. Training continued for a minimum of five sessions and until mice earned at least 30 pellets in three consecutive sessions. Following food self-administration, mice underwent jugular catheterization. As described previously^37^, isoflurane anesthesia was used, then the dorsal and ventral surfaces were shaved and disinfected. The right jugular vein was accessed via blunt dissection, and a sterile polyurethane catheter was inserted and secured to the vein. The catheter was externalized posterior to the scapulae using a felt mesh mount (Instech). Both dorsal and ventral incisions were sutured, and mice were pair-housed throughout the experiment, unless aggression or attrition necessitated individual housing. Postoperative care included analgesic treatment (Rimadyl 5 mg/kg) administered for three days. Mice were allowed to recover for 7 days before beginning intravenous self-administration. Catheter maintenance involved daily infusions of 0.05 mL gentamicin (0.33 mg/mL) and 0.05 mL heparinized saline (30 USP/mL) containing baytril (0.5 mg/kg). Patency was confirmed once per week using 0.05 mL brevital (3 mg/mL). Mice that did not exhibit loss of muscle tone were excluded from further testing.

After recovery, mice were trained to respond for cocaine (0.5 mg/kg/infusion, 30 μL delivered over 1.7 seconds) on a FR1 schedule using the inactive lever from prior food training. Cocaine infusions were administered via an armored tether connected to a swivel and syringe pump. Training sessions occurred six days per week, with the last day reserved for patency testing. Each drug infusion was paired with extinction of the house light, a compound cue (tone and stimulus light), and a 10-second timeout during which no further reinforcers were available. Sessions ended after acquiring 60 infusions or at 120 minutes. Acquisition was defined as self-administering ≥15 infusions across three sessions with a minimum 2:1 active-to-inactive lever press ratio, and completion of a minimum of 10 acquisition sessions. Other measures included drug intake (number of infusions), discrimination between active and inactive levers, and time to meet acquisition criteria.

Following acquisition, mice were tested on an FR1 schedule across two consecutive sessions for each of six descending doses of cocaine (1.0, 0.5, 0.25, 0.125, 0.063, and 0 mg/kg) for dose-response analysis. Mice were then returned to the baseline unit dose of 0.5 mg/kg for two sessions or until responding recovered and met acquisition criteria. Next, to assess motivation for cocaine in the same animals, three counterbalanced unit doses (1.0, 0.5, and 0.25 mg/kg) were tested across two consecutive sessions each, using a progressive ratio schedule in which each subsequent infusion required an increasing number of lever presses (1, 2, 4, 6, 9, 12, 16, 20…up to 240). Sessions ended after 4 hours or 1.5 hours without acquiring reinforcement. This test determines motivation, as reflected by the breakpoint ratio, which is the last ratio obtained for a dose of cocaine.

Mice resumed baseline self-administration (0.5 mg/kg/infusion) for two sessions before extinction training. During extinction, lever presses no longer resulted in any programmed outcome. Training continued for at least 10 days or until responding reached ≤30% peak active responding at 0.5 mg/kg/infusion. After extinction, sensitivity to cue-induced reinstatement was tested as a model for relapse-like behavior where presentations of previously cocaine-associated cues invigorate responding on the previously cocaine-reinforced lever. Mice were given a single non-contingent cue presentation, followed by cue presentations that were contingent on responses made on the previously active lever.

*Cohort 4: Food self-administration*

**Food self-administration**

Mice were initially restricted to 85% of their free-feeding weight. They were trained to lever press for chocolate-flavored food pellets (20 mg, grain-based precision pellets, Bio-Serv) delivered into the magazine of an operant conditioning chamber (Med-Associates). Responses to one lever were reinforced on a fixed ratio 1 (FR1) schedule, with a cue light illuminated above the lever for the entire session. Responses on the other lever were recorded, but had no programmed consequences. Sessions lasted 60 minutes or until 30 pellets were earned. Training continued for a minimum of five sessions, until mice earned 30 pellets for two consecutive sessions with a minimum 3:2 active-to-inactive lever press ratio on the second day. After a max of 10 total sessions, mice were tested on a FR3 schedule of reinforcement for 3 days. Next, reversal learning was assessed over 3 sessions by switching the contingencies of the lever. Here, previously inactive lever presses resulted in a reinforcer delivered on an FR3 schedule. Sessions ended after 30 minutes or after 30 pellets had been earned. Mice were next trained on a random interval (RI) schedule, first an RI30 schedule was used for 2 sessions, followed by an RI60 schedule for 3 sessions. Each session lasted 60 minutes or until 30 pellets were earned. Lastly, lever pressing was assessed under extinction conditions with no food pellets or cue light accessible. Sessions lasted 30 minutes.

*Cohort 5: Action-outcome contingency degradation*

**Action outcome contingency degradation**

A separate cohort of mice were trained to press two levers for food reinforcers under an FR1 schedule (5 sessions minimum), with a max of 30 reinforcers available on each lever. Sessions ended at 60 minutes or when mice acquired all 60 pellets. Mice were then trained for an additional 2 sessions at a random interval (RI) schedule 30 and then 3 sessions at RI60 to increase responding and bias towards habitual response strategies. Next, a modified version of action-outcome contingency degradation was used^36^. In the non-degraded session, one lever was retracted and the extended lever was reinforced using an FR1 schedule as during early training. In the degraded session, the opposite lever was available, and reinforcers were delivered at a rate matched to each animal’s reinforcement rate from the previous session. This response, therefore, becomes less predictive of reinforcement. These 2, 25-minute sessions were counter-balanced, as was the degraded contingency.

The following day, both levers were available during a 10-minute probe test under extinction conditions. The ratio for non-degraded/degraded lever presses was calculated. A goal-directed response strategy is shown by preferential engagement of the response that is likely to be reinforced (ratio over 1), while a habitual response strategy is to engage both familiar levers equally (ratio ~1).

We will only show data from the probe test and not lever pressing training, since this cohort consisted of mice with varied prior experience with the operant chamber (naïve or prior FR1/reversal training) and we have already shown findings for FR1 and RI training.

*Cohort 6:* *Cross fostering*

Pups from a control dam were fostered by a pCRD dam and vice versa. Mice in this cohort were evaluated for a subset of exploratory behavior such as open field and elevated plus maze, and reward-related behaviors such as conditioned place preference.

**Cross fostering**

Cross fostering occurred within 24 hours of birth. After recording the number of pups in each litter, all pups were removed and exchanged with a litter from the other group. For example, all control pups were transferred to a pCRD dam and all pCRD pups were transferred to the corresponding control dam. Mice were undisturbed throughout postnatal development until weaning.

**Blood collection**

We measured rhythms in corticosterone (CORT), as well as the glucocorticoid response to acute restraint stress. For CORT rhythms, adult mice (at least 12 weeks of age) exposed to pCRD, along with respective controls, were euthanized by cervical dislocations followed by rapid decapitation across time of day (TOD)(ZT2/7/10/14/19/22). Trunk blood was collected for measuring CORT rhythms. For the glucocorticoid response to acute restraint stress, adult pCRD and control mice were placed in 50-mL conical tubes between ZT2 and ZT7. Tail blood samples were obtained from a small incision at the tail tip at the beginning and end of restraint (0 and 15 minutes), as well as 15- and 105-minutes post restraint (30 and 120 minutes respectively).

All blood samples were collected in heparin-coated tubes (capillary and microcentrifuge) and kept on ice. Samples were centrifuged (1000 g for 10 min at 4 °C) to isolate the plasma and stored at -80 °C until analysis.

**Corticosterone ELISA**

Plasma corticosterone (CORT) samples were diluted 1:100 and CORT levels were quantified as instructed (ab108821, Abcam). Briefly, 25 µl of each standard and sample, followed by 25 µl of biotinylated CORT protein, were added to a 96-well plate and incubated for 2 hours at room temperature. After washing, 50 µl of streptavidin-peroxidase conjugate was added to each well and incubated for 30 min at room temperature. After washing, 50 µl of chromogen substrate was added to each well and incubated for 20 minutes in ambient light. Color developed inversely proportional to the amount of CORT in each well. The reaction was stopped with 50 µl of stop solution and the absorbance was immediately determined at a wavelength of 450 nm. CORT concentration was determined based on the standard curve. All samples were run in triplicate and mean values were used for data analysis.

**Statistical Analyses**

GraphPad Prism 10 and SPPS were used. Data are expressed as mean ± SEM with *p* ≤ 0.05 considered significant and 0.05 < *p* ≤ 0.1 considered trending. Measures collected in CRD dams and litters (e.g. sleep, pup mortality) were analyzed by two-tailed t-tests. Measures collected in adult offspring were analyzed by two-, three-, or four-way ANOVAs for effects of pCRD, sex and session/dose/lever if appropriate. ANOVAs with significant and trending interactions were followed by post-hoc tests with Bonferroni corrections for multiple comparisons where appropriate. If a significant interaction or effect of sex was found in a three-way ANOVA, data were separated by sex and two-way ANOVAs were performed. Since light and dark phase animals cannot be tested concurrently each phase was analyzed separately. For experiments assessing self-administration (drug or food) across RM (session, dose, or lever), data were analyzed by three-way ANOVAs for effects of pCRD, sex, and RM. Sex differences were commonly identified, therefore if effects of sex were found (main effects or interactions), data were separated by sex and analyzed by two-way ANOVAs to better examine effects across our RM. For sucrose preference, CPP and action outcome contingency degradation, one sample t-tests were used to determine whether significant preferences exist (compared to 50%, 0 and 1 respectively). Throughout, significant main effects and interactions are described. Outliers that were more than two standard deviations from the mean were excluded.

We quantified homecage activity using WakeActive and ActivityStatistics software, and sleep using SleepStats2p10 software (Signal Solutions, LLC, Lexington, KY, United States). Data were visualized as normalized actograms with 3 min bin sizes in ClockLab (Actimetrics, Wilmette, IL, United States).
